# A benchmarking workflow for assessing the reliability of batch correction methods

**DOI:** 10.1101/2025.08.01.668073

**Authors:** Elfried Salanon, Blandine Comte, Delphine Centeno, Stéphanie Durand, Estelle Pujos-Guillot, Julien Boccard

## Abstract

One of the most pervasive challenges in large-scale untargeted metabolomics is short and long-term analytical variability introducing the necessity of batch effect correction. In this context, several strategies and methods have been developed to limit those effects, either by monitoring the data generation process to maximize reproducibility or by applying post-analysis data correction. Different evaluation frameworks, either assessing the degree of bias in the data through visual tools or quantitative indicators, or evaluating the prediction performance of known biomarkers, were also proposed. However, there is currently no clear consensus on how to evaluate batch correction methods. This work offers a strategy to assess multiple dimensions of batch correction efficiency within a comprehensive and reliable framework, designed to assess the effectiveness and reliability of batch correction methods. Based on Mahalanobis Conformity Index (MCI), it provides a multivariate and covariance-aware metric to quantify within- and between-batch variability. Additionally, it combines visualization techniques (Principal Component Analysis (PCA) and Multivariate INTegrative (MINT) PCA) with numerical indicators (batch dispersion, Coefficient of Variation), supporting both multidimensional and metabolite-specific evaluations. Lastly, this novel approach integrates statistical tools alongside chemistry-based metrics for method overfitting and overcorrection assessment. Applied within a use case for comparing LOESS-based and ComBat correction methods, the present workflow provided a structured approach to systematically assess the reliability of batch corrections, ensuring both data intercomparability and biological relevance in metabolomics studies.

**Author summary:** The assessment of batch correction is a challenge in metabolomics, where there is no consensus for a define strategy, making it a complex task. However, knowing the impact of batch correction on the datasets and consecutive possible impact on downstream statistical analyses, providing a reliable framework for its assessment is a cornerstone for reproducible results. The objective of the present work was to provide a framework and a set of interpretation tools combining numerical indicators, as well as diagnostic plots, for assessing the reliability of batch correction methods. We introduced a robust evaluation framework centered on the Mahalanobis Conformity Index, providing a multivariate and covariance-aware metric to quantify within- and between-batch variability. By coupling this index with visual tools (based on Principal Component Analysis (PCA) and Multivariate INTegrative (MINT) PCA), as well as compound-level diagnostics, we enabled a fine-grained and interpretable comparison of correction strategies, highlighting their strengths and potential pitfalls.

## Introduction

Recent advances in analytical techniques, particularly in mass spectrometry (MS) allowed increasing data quality in metabolomics in terms of sensitivity and robustness, opening the door to its large-scale application (1). However, in MS-based untargeted metabolomics, even if the multiple sources of random error are generally reduced through good laboratory practices, systematic errors, often caused by impaired analytical systems, may affect the quality of collected data. As untargeted metabolomics does not involve absolute quantification, this can hinder the discovery of biomarkers by increasing the risk of false positives or negatives (2). These batch effects are usually evaluated *via* repeated measurements of samples using internal standards or certified reference materials (3).

In this context, several strategies and methods have been developed to limit batch effects (4–8). Some proposed quality controls strategies consist in monitoring the data generation process to maximize reproducibility. Other alternative strategies are based on post-analysis data correction. In that case, several methods rely on the use of quality control (QC) samples to estimate bias and evaluate a correction function, as the LOcally Estimated Scatterplot Smoothing (LOESS) approaches or the linear model-based correction (Dunn et al., 2011). Alternatively, Giodan et al (9) proposed to apply data driven strategies independent of QC samples, such as ComBat developed by Johnson et al (10) and BER (batch effect removal)(9).

There is currently no clear consensus on how to evaluate batch correction methods. Researchers proposing new approaches or assessing existing strategies for routine use generally rely on different categories of evaluation frameworks (9,10). The first type of strategy aims to assess the degree of bias in the data, either through visual tools such as Principal Component Analysis (PCA), and the explained variance, t-distributed stochastic neighbor embedding (t-SNE), relative expression plots (11,12), or through quantitative indicators. These include the D-ratio, coefficient of variation (CV), or correlation coefficients observed in diluted QC samples (3,13). A second evaluation strategy is based on the prediction performance of known biomarkers identified in previous studies. In such cases, a panel of established metabolites is used to assess whether their discriminatory power is preserved or improved after applying a given batch correction method, using previously published datasets (9).

However, although these methods coexist, no standardized workflow/tool is currently available for the comprehensive assessment of either newly developed correction algorithms, or of the performance of those used in routine applications. Furthermore, because batch effects arise from both intra- and inter-batch variation as well as study- and platform-specific artifacts (3,13), their evaluation cannot rely on a single indicator. While multidimensional visualization tools such as PCA provide a global view of performance, they should be complemented by univariate assessments that focus on specific metabolites. Metrics such as CV, intra-group dispersion, or D-ratio offer this metabolite-level perspective. It is also important to assess how well the correction addresses intra- and inter-batch variation (3,14). Finally, while most current approaches rely solely on statistical indicators, a complete workflow should also include evaluations from chemical and biological standpoints.

The workflow and tools presented in this work offer a strategy to assess multiple dimensions of batch correction performance (Figure 1). They combine visualization techniques with numerical indicators, supporting both multidimensional and metabolite-specific evaluations. Additionally, the approach integrates statistical tools alongside chemistry-based metrics such as isotopic ratio consistency providing a more comprehensive and reliable framework for method overfitting and overcorrection assessment.

**Figure 1:**
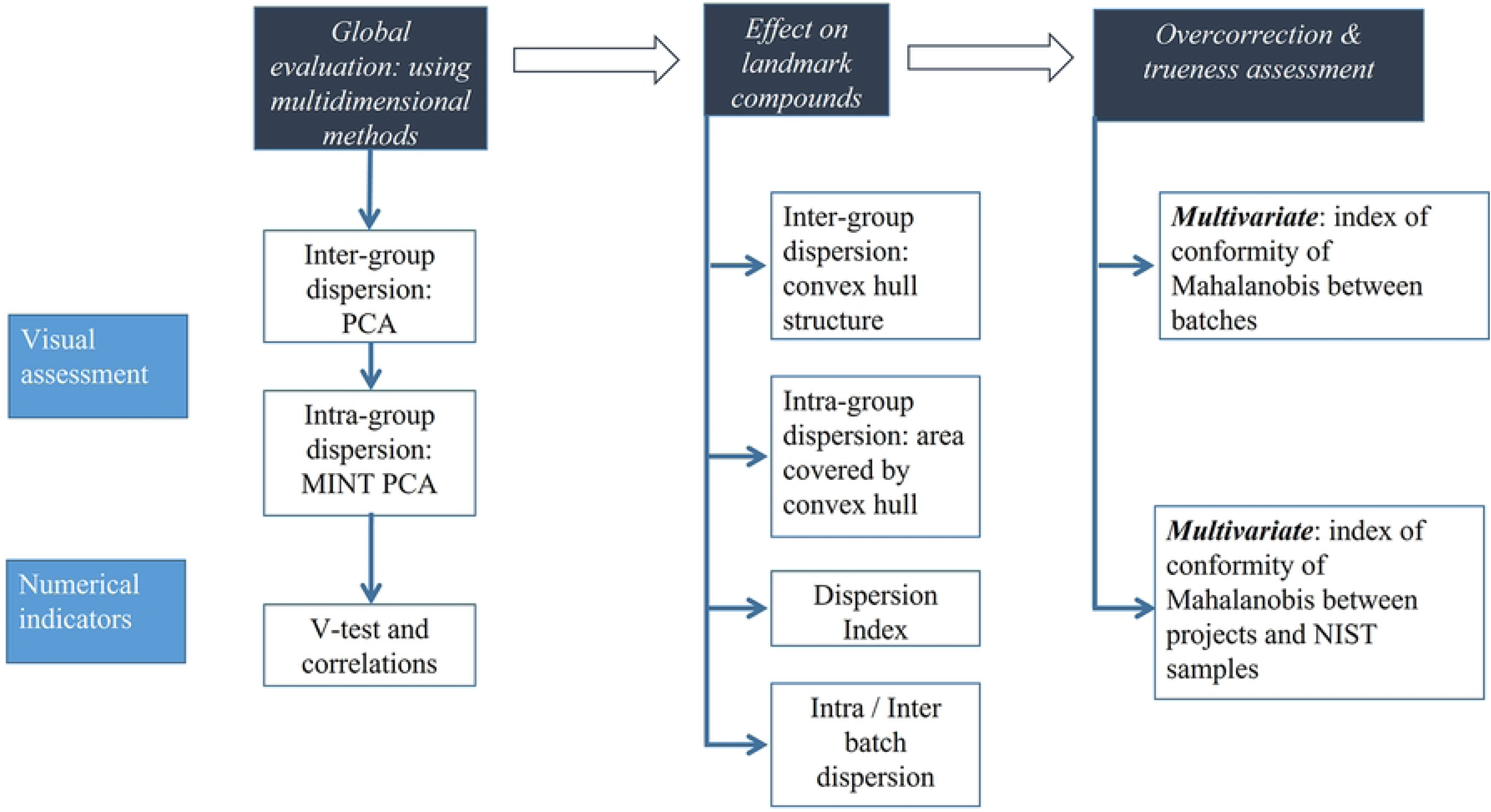
Comprehensive workflow for assessing the reliability of batch correction methods

## 2. Results & Discussion

### 2.1. Multivariate assessment

#### ➢ Inter-batch assessment

##### • Visual evaluation

In the Figure 2a, the QCs linked with different batches appeared to be clearly separated on the PCA score plot (PC1 *vs.* PC2) before batch effect correction. Batch QCs were well centered after the LOESS correction as presented in Figure 2b and the same trends were observed with the ComBat-based correction (Figure 2c). However, a less uniform distribution was obtained with the LOESS method. Moreover, the correlation of the Batch and the injection order *(r = 0.61 before correction vs. -0.02 after LOESS” & -0.01 after “ComBat”)*, which were important on the first two principal components, could not be observed anymore after correction with both methods. This was followed by a reduction of the proportion of variance explained by those two dimensions *(Dimension 1 = 21.36% before correction vs. 15.05% after “LOESS” & 17.39% after “ComBat”; Dimension 2 = 13.52% before correction vs. 4.33% after “LOESS” & 4.28% after “ComBat”)*. This showed a global improvement in the data structure by reducing the inter-batch distance. However, knowing that PCA offers a relative evaluation of the variability contained in the data, it should be considered as multidimensional assessment and a first exploration that needs to be completed with other criteria.

**Figure 2:**
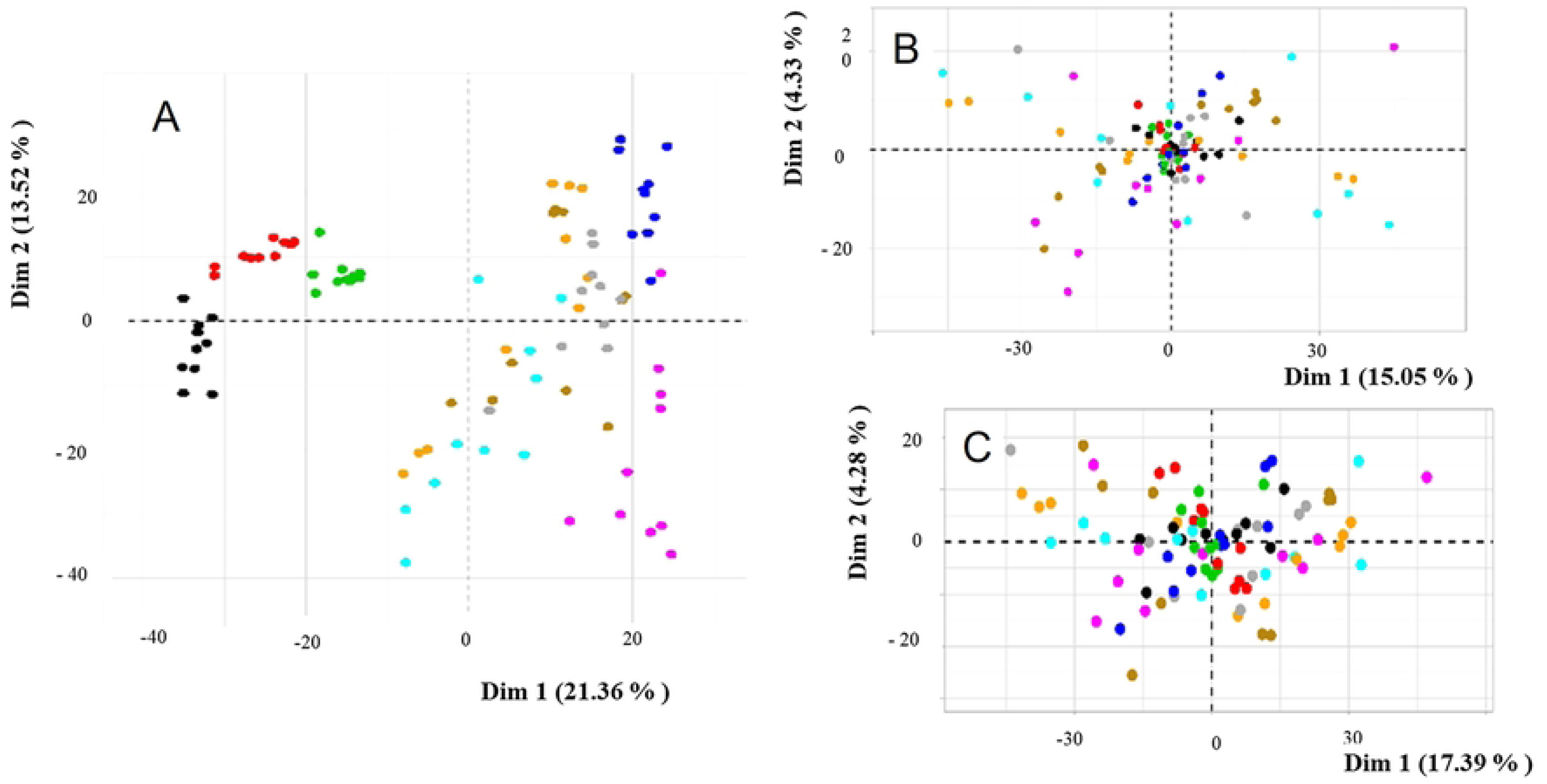
PCA score plot using the QC samples analyzed across nine batches on a large-scale study. a: Without batch correction, b: After application of the LOESS batch correction, c: After application of the ComBat correction on the data. The different colors represent the different analytical batches

##### • Clustering

Figure 3 presents the clustering using hierarchical cluster analysis after PCA using 5 components. As presented in figure 3a, before correction, cluster 1 was associated with batch 1 *(Value-test (V-test) =7.37),* while batches 2 & 3 were linked to cluster 2 *(V-test = 5.53).* Cluster 3 was composed of batches 9 & 8 with a v-test of 2.17 and the clusters 5 & 6 included respectively batch 5 *(V-test = 6.38)* & batch 4 *(V-test = 7.37).* Three clusters were identified after LOESS correction (figure 3b). The first cluster was linked to batch 8 *(V-test =2.09).* Batch 9 was associated with cluster 3 (*V-test: 3.02*). The other batches were correlated to cluster 2. The ComBat-based batch correction revealed three clusters, with batch 5 being the only associated to cluster 3; all the others clusters were found to be a mix of all the batches. This analysis revealed a reduction of the batch effect, with an improvement of the homogeneity of the clusters. However, some trends could still be observed for batches 8 and 9 after applying the LOESS methods, and batch 5 after the ComBat method. This could raise some questions about the specificities of those batches in order to identify any potential event that could have occurred during the data generation process. While classical PCA was limited to a visual assessment of the inter-batch changes before and after correction, clustering allowed groupings to be more objectively evaluated with respect to batch effect or to a potential biological trend. This could be investigated via the membership of the different batches to the clusters, highlighting others sources of effect, which may be of interest.

**Figure 3:**
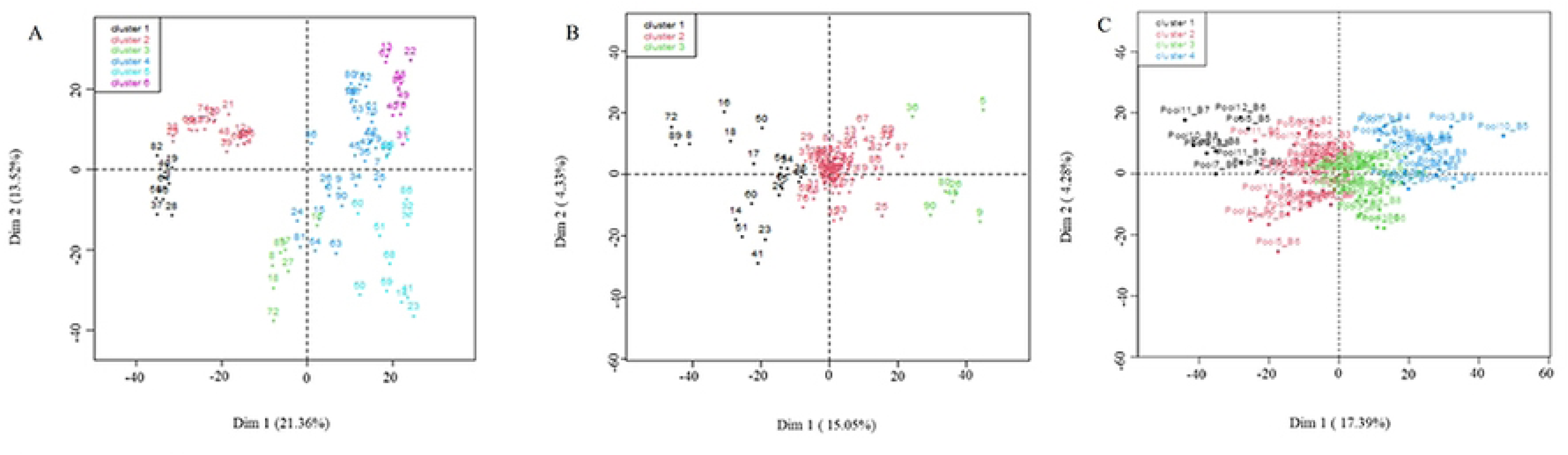
Projection in two dimensions of the results of the Hierarchical Cluster Analysis, using QC samples analyzed within the nine batches. ***a****: Without batch correction, **b**: After LOESS correction, **c**: After ComBat correction. The different colors represent the different batches*

##### • Numerical evaluation

Table 1 presents MCI values before and after batch correction calculated on QC samples. The inter-batch correction corresponds to the multivariate assessment of the distance of the barycenter (median) of each batch to the first. Before correction, the median distance was 480.47, while MCI values of 16.53 and 16.63 were observed after correction by LOESS and ComBat methods, respectively. This measurement gives an objective evaluation of the multivariate distance (>1,800 features) of the different batches to the first. This value is meant for comparison relatively to the value before correction. In this use case, there was an improvement of the global inter-batch comparability shown by the decrease of the distance between the different batches to the reference. This is an expected behavior of the correction methods aiming to reduce the difference between the barycenter of the different batches. Moreover, the comparison of the two batch correction methods revealed that they globally performed equally. This is consistent with the results of the visual assessment of the inter-batch variation, showing that both correction methods reduced significantly the inter-batch effect.

**Table 1:** Inter-batch MCI, before and after correction, using the 100 QC samples analyzed within nine batches on a large-scale study.

| Batch | Before batch correction | LOESS Batch correction | ComBat batch correction |
| --- | --- | --- | --- |
| Batch2 | 436.23 | 15.56 | 13.98 |
| Batch3 | 521.44 | 15.92 | 15.97 |
| Batch4 | 477.64 | 16.62 | 16.21 |
| Batch5 | 456.66 | 23.59 | 17.47 |
| Batch6 | 526.62 | 16.44 | 16.91 |
| Batch7 | 531.08 | 15.98 | 16.35 |
| Batch8 | 414.25 | 19.96 | 21.20 |
| Batch9 | 483.30 | 20.18 | 18.71 |
| Inter-batch MCI | 480.48 | 16.53 | 16.63 |

#### ➢ Intra-batch assessment

Figure 4 presents the multivariate evaluation of the intra-batch dispersion using MINT PCA. Only little improvement was observed after correction, but LOESS seemed to generate more spread as presented in Figure 4b. Numerical estimators were further investigated to better evaluate the dispersion and complement this first visual assessment.

**Figure 4:**
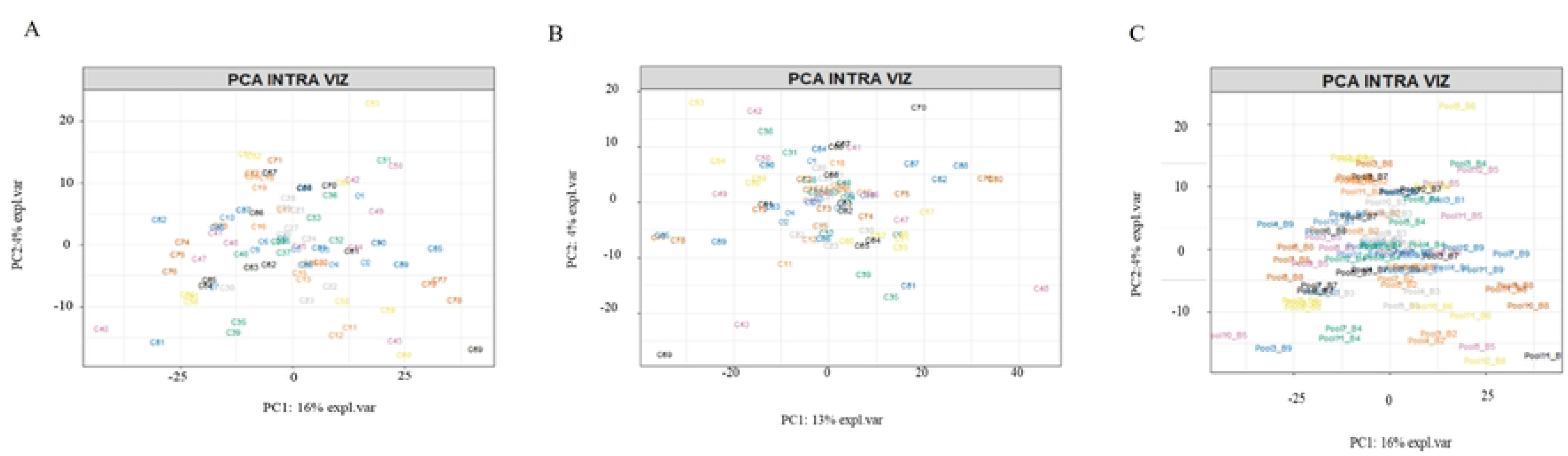
MINT PCA score plots using QC samples analyzed within the nine batches on a large-scale study. **a**: Without batch correction, **b**: After LOESS batch correction and **c**: After ComBat correction. The different colors represent the different batches

As presented in table 2, the multivariate assessment of the intra-batch dispersion using the median MCI to their barycenter implying all the variables (> 1,800 features), revealed little differences between the batches except for batch 9, which had a particularly low value of intra-batch MCI. The LOESS correction showed a lower intra-batch MCI, while the ComBat correction method did not seem to provide any major improvement. The expected behavior for the intra-batch assessment is a reduction of MCI in both the different batches and the global intra-batch value. This decrease revealed a multivariate improvement of the distance of each QC sample to the barycenter of its batch, showing an enhancement in the batch wise harmonization.

**Table 2:** Intra-batch MCI, before and after correction, using the QCs analyzed within nine batches on a large-scale study.

| Batch | Before<br>correction | LOESS correction | ComBat<br>correction |
| --- | --- | --- | --- |
| Batch1 | 8.25 | 8.23 | 8.27 |
| Batch2 | 8.22 | 7.96 | 8.10 |
| Batch3 | 8.10 | 7.49 | 8.12 |
| Batch4 | 8.21 | 7.24 | 8.21 |
| Batch5 | 8.25 | 7.76 | 8.37 |
| Batch6 | 8.39 | 7.99 | 8.37 |
| Batch7 | 8.68 | 6.85 | 8.67 |
| Batch8 | 8.22 | 7.06 | 8.24 |
| Batch9 | 7.06 | 7.19 | 8.64 |
| Global Intra-batch<br>MCI | 8.22 | 7.49 | 8.27 |

Moreover, analyzing the shapes of convex hulls of the different batches in Figure 5 revealed that ComBat preserved the shape of the dispersion before and after correction, while the LOESS correction was more prone to change it. This conservative behavior of interest for any biological information could be explained by the inner structure of ComBat, which is based on means and scale (Johnson et al., 2012), while the LOESS method is based on the QCs to fit a correction function via a local regression model. However, this conservative behavior is not what is expected when considering batch correction based on repeated QCs. This assessment is multivariate and is intended to give a global idea of the improvement of the quality. However, targeted evaluation using individual metabolites, *i.e.* landmark compounds, constitutes an interesting complement to evaluate some specific aspects in a univariate way.

**Figure 5:**
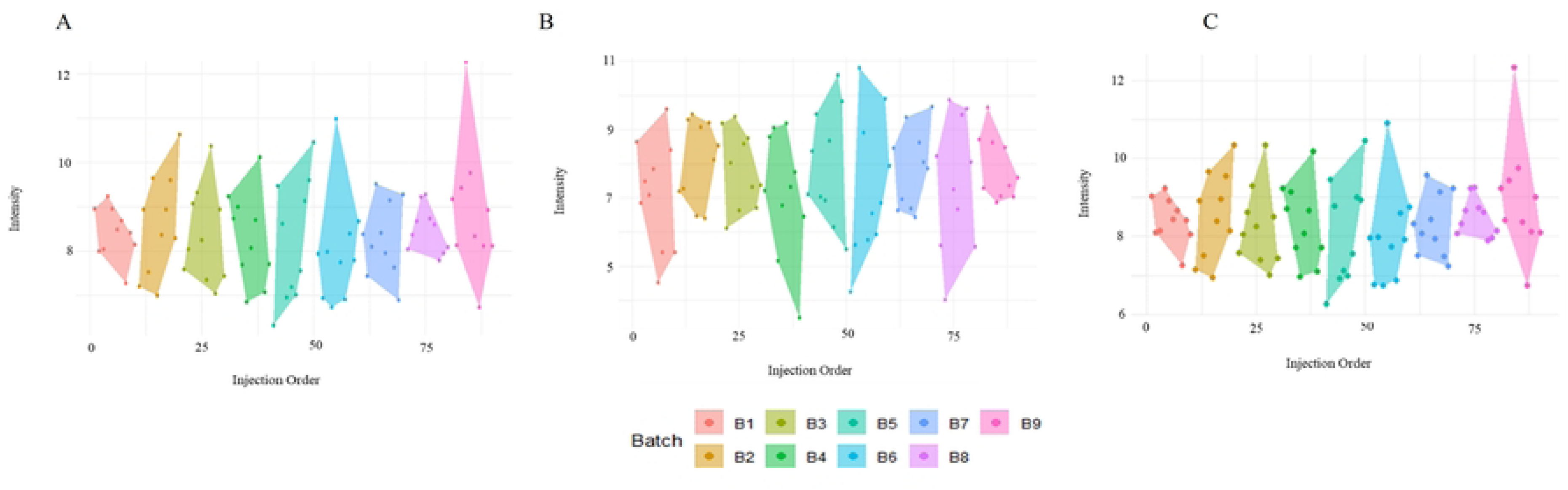
Intra batch MCI convex hulls, before and after correction, using QC samples analyzed within the nine batches. **a**: Without batch correction, **b**: After LOESS correction and **c**: After ComBat correction. The different colors represent different batches

### 2.2 Univariate assessment using landmark compounds

#### ➢ Inter-batch effect

Table 3 presents the coefficients of variation as well as the intergroup dispersion of the selected landmark compounds. Results showed a significant decrease in those indicators after correction using both correction methods. Based on the coefficient of variation, LOESS outperformed ComBat except for one amino-acid (L-threonine). This is consistent with the results of the inter group dispersion which revealed the same trends for most indicators. These results are also consistent with the MCI numeric assessment in a multivariate context, showing that LOESS outperformed ComBat for inter-batch correction. Those results are confirmed by the visual assessment of landmark compounds bivariate dispersions. Indeed, while ComBat and LOESS aim to correct for inter-batch differences, ComBat tends to increase batch-wise dispersion in order to get an alignment of the mean intensity of the different batches, while the intra-batch dispersion was still very large in the different batches.

**Table 3:** Inter-group dispersion and coefficients of variation of representative landmark compounds, using the QCs analyzed within nine batches on a large-scale study of more than 992 participants.

| Metabolites | Inter-batch dispersion |  |  | Coefficient of variation |  |  |
| --- | --- | --- | --- | --- | --- | --- |
|  | Before correction | LOESS | ComBat | Before correction | LOESS | ComBat |
| Caffeine | 22826.1 | 6744.8 | 6233.1 | 16.7 | 9.9 | 9.6 |
| Creatine | 28386 | 5606 | 5211.1 | 30.7 | 11.3 | 14.1 |
| Creatinine | 10916.4 | 3305.7 | 5138.3 | 20.4 | 8.8 | 9.4 |
| Decanoyl-L-carnitine | 2194.3 | 231.2 | 398.2 | 21.9 | 12.8 | 15.2 |
| Hexanoyl-L-carnitine | 950.1 | 64.4 | 109.7 | 21 | 14.8 | 15 |
| Isovaleryl-L-carnitine | 672.4 | 37.2 | 72.3 | 16 | 9.9 | 10.8 |
| L-Glutamic acid | 3865.7 | 514.9 | 1030.2 | 21.2 | 7.3 | 9.3 |
| L-Histidine | 865.3 | 386.2 | 155.2 | 10.3 | 6 | 6.4 |
| L-Leucine | 37435.1 | 6328.5 | 44886.9 | 19.6 | 11.1 | 14.8 |
| L-Lysine | 68703.2 | 132214 | 56210.7 | 28.7 | 15.1 | 15.7 |
| L-Methionine | 36473.2 | 36443.8 | 525 | 24.1 | 19.6 | 23.5 |
| L-Threonine | 709.2 | 172.9 | 4268.2 | 37.9 | 17.5 | 6.9 |
| L-Tyrosine | 25433.5 | 19904.5 | 10364.8 | 19 | 11.4 | 11.4 |
| L-Valine | 214459.9 | 32116.3 | 68083.5 | 12.5 | 7.8 | 8.5 |
| L-Tryptophan | 28782.6 | 3290.9 | 13383.6 | 13.8 | 8.3 | 9.6 |
| Lyso PC(P-16:0) | 5353.8 | 546.8 | 948.8 | 34.6 | 19.8 | 23.5 |
| Myristoyl-L-carnitine | 525.6 | 26.9 | 31.1 | 27.1 | 15.1 | 17.8 |
| Phenylacetyl-L-glutamine | 5926.4 | 149.8 | 330.7 | 27.9 | 9.7 | 11.8 |
| Theophylline | 4496.3 | 815 | 744.7 | 11.4 | 5.5 | 6.6 |
| Uric acid | 50902.4 | 45419.5 | 12975.6 | 18.2 | 17 | 12.1 |

#### ➢ Intra-batch correction

Table 4 presents the intra group dispersion and the dispersion index of the landmark compounds, showing that intra group dispersion was globally decreased after correction. However, it should be noted that these values increased after correction for some of the landmark compounds. Moreover, intra group dispersion was globally lower after LOESS correction compared to Combat. These findings are consistent with the results of the multivariate assessment regarding intra group dispersion. The expected behavior is a decrease in the value of the intra group dispersion showing an improvement. However, the increase in the dispersion index for most of the landmark compounds indicates that both correction methods led to better results in the correction of the intergroup effect than in the intra group effect correction. Indeed, the intra-batch variability is more strongly connected to the structure of the analytical shift resulting from the analytical sequence, while the inter-batch effect is more related to global differences, making it more straightforward to correct.

**Table 4:** Intra-group MCI, before and after correction, using the QCs analyzed within nine batches on a large-scale study of more than 992 participants.

| Metabolites | Intra group dispersion |  |  | Dispersion Index |  |  |
| --- | --- | --- | --- | --- | --- | --- |
|  | Before correction | LOESS | ComBat | Before correction | LOESS | ComBat |
| Caffeine | 45772.0 | 25869.4 | 37157.7 | 2.0 | 3.8 | 6.0 |
| Creatine | 91456.8 | 37553.6 | 82437.1 | 3.2 | 6.7 | 15.8 |
| Creatinine | 88327.6 | 39434.3 | 86291.9 | 8.1 | 11.9 | 16.8 |
| Decanoyl L-carnitine | 8693.6 | 5159.8 | 8400.3 | 4.0 | 22.2 | 21.0 |
| Hexanoyl L-carnitine | 1956.0 | 1763.4 | 2144.0 | 2.1 | 26.9 | 19.4 |
| Isovaleryl-L-carnitine | 1448.0 | 932.9 | 1267.6 | 2.2 | 24.4 | 17.3 |
| L-Glutamic acid | 6540.8 | 4539.1 | 6973.1 | 1.7 | 8.8 | 6.8 |
| L-Histidine | 2889.6 | 2246.1 | 2655.1 | 3.3 | 5.8 | 17.0 |
| L-Leucine | 57322.4 | 48727.2 | 58905.8 | 1.5 | 7.7 | 1.3 |
| L- Lysine | 57539.6 | 52685.1 | 110698.4 | 0.8 | 0.4 | 2.0 |
| L Methionine | 72570.8 | 67882.9 | 4657.5 | 2.0 | 1.9 | 8.9 |
| L Threonine | 2815.2 | 2841.9 | 91934.3 | 4.0 | 16.3 | 21.5 |
| L-Tyrosine | 111783.2 | 105240.4 | 119668.1 | 4.4 | 5.3 | 11.5 |
| L-Valine | 969595.6 | 1038837.6 | 1046562.6 | 4.5 | 32.3 | 15.4 |
| L-Tryptophan | 55238.0 | 50252.1 | 51544.6 | 1.9 | 15.3 | 3.9 |
| Lyso PC(P-16:0/0:0) | 6618.0 | 6769.8 | 8337.1 | 1.2 | 12.4 | 8.8 |
| Myristoyl L-carnitine | 1400.4 | 1110.8 | 1490.7 | 2.7 | 39.8 | 46.5 |
| Phenylacetyl L-glutamine | 9936.0 | 4056.6 | 2871.0 | 1.7 | 26.9 | 25.4 |
| Theophylline | 6526.0 | 4750.7 | 7658.8 | 1.5 | 5.8 | 25.4 |
| Uric acid | 51623.6 | 54222.6 | 72398.5 | 1.0 | 1.2 | 10.3 |

These findings were confirmed by the visual representations of the bivariate dispersion of landmark compounds in Supplemental Figure 1. While there was a decrease in the inter-batch distance, the intra-batch dispersion of some batches increased in order to have a global alignment. For example, the intra-batch variability of uric acid was preserved without improvement or deterioration after LOESS correction, while increased intra-batch dispersion was observed after ComBat. The same trends were observed for lysine and L-methionine.

#### ➢ Effect of the correction on the isotopic ratios

Table 5 presents the isotopic ratios of several landmark metabolites before and after correction using LOESS and ComBat methods, considering only measurements above the S/N >10 threshold. The goal was to assess how batch correction algorithms preserve chemically known value, using theoretical natural isotope ratios as reference values.

**Table 5:** Isotopic ratios before and after correction, using the QC samples analyzed within nine batches.

| Landmark compounds | Average intensities |  |  | Isotopic Ratio |  |  |  | Standard deviation of the Isotopic ratio |  |  |
| --- | --- | --- | --- | --- | --- | --- | --- | --- | --- | --- |
|  | Without | LOESS | ComBat | Without | LOESS | ComBat | Theoretical | Without | LOESS | ComBat |
| L-Arginine | 255431.07 | 252915.41 | 255280.12 | 8.65 | 8.69 | 8.55 | 8.20 | 1.10 | 1.10 | 1.23 |
| Hippuric acid | 124778.80 | 125936.42 | 125081.02 | 11.99 | 11.83 | 11.97 | 10.30 | 1.60 | 1.41 | 1.32 |
| Theobromine | 175638.69 | 175717.00 | 175633.78 | 9.44 | 9.45 | 9.21 | 9.20 | 1.57 | 1.55 | 1.82 |
| L-Tyrosine | 1648241.47 | 1648516.32 | 1647534.54 | 11.49 | 11.46 | 11.49 | 10.30 | 0.42 | 0.49 | 0.40 |
| Caffeine | 594063.33 | 599396.73 | 593703.24 | 10.85 | 10.86 | 10.85 | 10.30 | 0.78 | 0.69 | 0.65 |
| L-Tryptophan | 2467803.36 | 2463565.55 | 2463281.09 | 14.55 | 14.61 | 14.56 | 12.80 | 1.53 | 1.33 | 0.67 |
| Phenylacetylglutamine | 109072.62 | 109755.39 | 109691.28 | 16.41 | 16.08 | 16.19 | 15.10 | 2.52 | 2.20 | 2.51 |
| Decanoyl-carnitine | 79347.80 | 79185.98 | 79435.94 | 19.61 | 19.67 | 19.62 | 19.30 | 2.06 | 1.90 | 2.12 |
| Creatinine | 1326372.13 | 1329412.24 | 5025833.73 | **5.34** | **5.38** | **6.85** | **5.50** | 1.16 | 1.16 | 1.15 |
| L-Proline | 5016879.31 | 5057731.28 | 977707.24 | **6.85** | **6.86** | **7.13** | **6.00** | 0.25 | 0.18 | 0.21 |
| Pipecolic acid | 978320.80 | 982817.45 | 875383.14 | **7.30** | **7.31** | **5.22** | **7.10** | 2.79 | 2.27 | 1.59 |
| Creatine | 873001.58 | 878834.40 | 461399.23 | **5.46** | **5.50** | **8.13** | **5.60** | 1.06 | 1.27 | 1.40 |
| L-Leucine | 471825.89 | 472451.13 | 150803.45 | 7.95 | 7.88 | 7.19 | 7.10 | 1.24 | 1.02 | 1.77 |
| Trigonelline | 150620.11 | 151560.06 | 715147.65 | 7.64 | 7.45 | 7.30 | 8.10 | 1.37 | 1.93 | 1.47 |
| L-Lysine | 724408.40 | 722198.92 | 1196698.18 | 7.23 | 7.19 | 7.07 | 7.50 | 1.69 | 1.58 | 1.87 |
| L-Glutamic acid | 129587.76 | 130848.18 | 129510.87 | **6.01** | **6.01** | **5.74** | **6.00** | 1.51 | 2.63 | 1.38 |
| L-Methionine | 1198202.47 | 1201474.60 | 1196698.18 | 7.06 | 7.06 | 7.07 | **6.80** | 50.61 | 35.85 | 3.00 |

\*numbers in bold are metabolites where more important differences are found in the estimations.

Both LOESS and ComBat yielded modest improvements in isotopic ratio accuracy compared to uncorrected data. LOESS outperformed ComBat in 12 out of 17 compounds in terms of absolute deviation from the theoretical value, while ComBat showed improvement in 10 compounds. However, ComBat introduced significant overcorrections in several cases (e.g., Pipecolic acid, Creatine), where the isotopic ratio deviated further from the theoretical reference than in the raw data.

To further evaluate the correction performance, we analyzed both absolute differences (in ratio units) and relative errors (percentage deviation from theoretical values). In most cases, the two criteria were concordant; meaning compounds with high absolute deviations also exhibited high relative errors. However, a few compounds with low theoretical ratios (e.g., Creatinine or L-Proline) showed small absolute deviations but relatively high relative errors, revealing that relying solely on absolute differences may underestimate the extent of correction bias in low-abundance contexts. Thus, both criteria are complementary and should be jointly considered to avoid misinterpretation.

This analysis highlights that, isotopic ratios when theoretical expectations are known, can serve as internal quality control metrics for batch effect correction pipelines. They offer a chemical anchor point that allows researchers to assess not only the agreement of corrected values but also to detect overcorrection artifacts, as illustrated by ComBat in some cases. For instance, ComBat produced isotopic ratios up to 1.9 units away from theoretical values in compounds where the uncorrected data was already close to the theoretical value.

Although both correction strategies offered minor improvements, LOESS correction yielded better overall consistency and preserved chemical consistency more reliably. The convergence between absolute and relative criteria supports its robustness. The use of known isotopic ratios should be encouraged as a practical, compound-level tool for validating and benchmarking batch correction methods in untargeted metabolomics workflows.

## 3. MATERIALS and METHODS

### 3.1. Preliminary method development: the Mahalanobis Conformity Index

The Mahalanobis Conformity Index (MCI) is the weighted median of the Mahalanobis distance(15) (Mahalanobis et al., 1933) used to assess the concordance of observations within a given group relative to a reference, taking into account the covariance structure of the data. This measure, providing a multivariate indicator, is particularly relevant for assessing batch effect correction in metabolomics and other fields requiring data harmonization, where variations introduced by experimental conditions must be minimized, while preserving meaningful biological differences.

#### ➢ Definition and Mathematical Formulation

The MCI is defined as a weighted average or median of Mahalanobis distances, capturing the dispersion of observations around a reference point. Mathematically, it is expressed as:

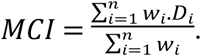

Where 𝐷_𝑖_ represents the Mahalanobis distance of observation 𝑖 to the barycenter, defined as the median of the reference group. The weighting factor 𝑤_𝑖_allows for flexible adjustments, incorporating measurement reliability or other relevant features. The Mahalanobis distance itself is computed as (Mahalanobis et al., 1933):

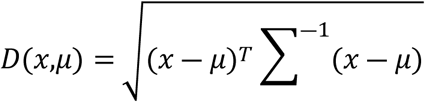

Where 𝑥, is an observation, 𝜇 is the barycenter, and 𝛴 is the covariance matrix, which can be adapted to different assumptions about the data structure. Unlike conventional distance metrics, this formulation accounts for the correlation between variables, making it well suited for high-dimensional, structured datasets such as those collected in metabolomics.

#### ➢ Properties and justification

A key MCI property is its invariance to linear transformations, inherited from the Mahalanobis distance. This ensures that comparisons between datasets remain meaningful even when scaling or affine transformations are applied. Additionally, MCI is sensitive to covariance variations, allowing it to capture dependencies between variables that might otherwise be overlooked with simpler distance metrics. The measure is also stable in the presence of extreme values, particularly in its robust version, which uses the median as a reference and applies shrinkage-based or robust estimations of the covariance matrix to mitigate the influence of outliers.

#### ➢ Versions of the MCI

MCI exists in two main versions: a standard and a robust extension. The standard version leverages the classical covariance matrix to calculate Mahalanobis distances, making it more responsive to fine-scale variations in the data. This approach is particularly useful when batch effects are subtle and need precise evaluation. However, this sensitivity also makes it more vulnerable to outliers and heterogeneity. The robust version, by contrast, employs alternative methods to enhance stability. Instead of using the mean as a reference, it adopts the multivariate median, ensuring that extreme values do not disproportionately influence the measure. Furthermore, when the covariance matrix is ill-conditioned or singular, shrinkage techniques are applied to ensure reliable inversion. This makes the robust version more suitable for datasets where batch effects are severe or where variability within batches is highly heterogeneous.

### 3.2. Proposed workflow for the batch effect assessment

The proposed evaluation of batch effect is a multistep approach, combining multidimensional and univariate tools. While the multidimensional tools provide a global view of the correction performance, univariate evaluation using landmark compounds provides specific insights as these compounds act as diagnostic features of the correction effect covering the diversity in terms of retention time.

*Step 1: Global evaluation: using multidimensional methods* Concerning the multidimensional assessment, Principal Component Analysis (PCA) (Jollife et al., 2016) and the Multivariate INTegrative (MINT) PCA (Rohart et al. 2016; 2017) are respectively used to visualize inter and intra-batch effects. Moreover, using the barycenter of each batch as reference, the MCI of the different measurement points provides a multidimensional numeric assessment of the intra-batch effect, while considering the median of batch values provide a global assessment. Concerning the inter-batch assessment; we use the MCI of the barycenter of the different batches, to the first batch as reference. As proposed recently, Mahalanobis distances of samples to their batch barycenter can be efficiently visualized using convex hulls (Salanon et al., 2024). It provides a simple graphical evaluation of the global dispersion within batch before and after correction.
*Step 2: Targeted assessment on landmark compounds (See appendix 2 for the selection process of the targeted landmark compound)* Prior choice of the landmark compounds must be done based on expert knowledge to cover the range of physicochemical diversity of the detected metabolites. Based on these landmark compounds, univariate assessment aims at evaluating the effect of the correction in specific areas. The coefficient of variation and the inter group dispersion (Van Der Kloet, 2009; Salanon et al., 2024) are used for numeric assessment of the inter-batch correction, while the inter group dispersion and the dispersion index provide a numeric estimation of the inter-batch effect (Salanon et al., 2024). A visual evaluation can also be performed using the convex hull based bivariate dispersion as recently proposed by Salanon et al., 2024. It allows displaying the correction effect on the inter and intra-batch variability *via* the landmark compounds in a reliable way.
*Step 3: Overcorrection & agreement with theoretical isotopic values assessment* In the context of evaluating batch correction methods, it is essential not only to quantify how well they reduce batch-related variability, but also to verify that they do not overcorrect the data. One way to monitor this is by using the Multivariate Correction Index (MCI), calculated as the distance between QC samples and a reference material (e.g., NIST SRM 1950 for plasma). The expected behavior after correction is a decrease in MCI between QCs across batches, while maintaining (or even increasing) the distance between the average QC signal and the reference material. In short, the correction should bring QCs together, but not distort the biological information present in the data.

To further evaluate potential overcorrection, we used the natural isotopic ratios of a set of landmark compounds, which have known theoretical values. These serve as internal standards to assess how much the correction respects true chemical information. For each compound, we computed the absolute error and the relative error as follows:

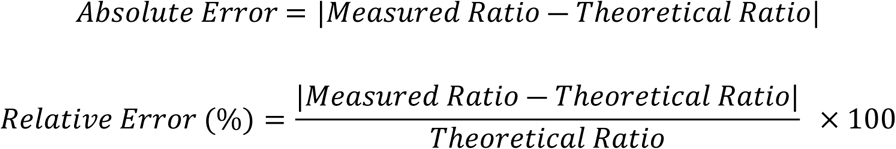

Using both metrics is key: absolute error captures how far we are from the theoretical value in raw units, while relative error flags when a small-looking deviation is actually large in proportion to the expected ratio (which matters especially for low-abundance compounds). As a rule of thumb, we considered that a relative error > 5% indicates a potentially meaningful deviation. These thresholds are consistent with typical tolerances in isotopic quantification in biological matrices for metabolomics, using such a QToF instrument. By combining this approach with MCI, we can detect whether correction methods preserve the chemical information or, on the contrary, introduce distortions. In that sense, isotopic ratios are more than a validation step: they offer a practical, compound-level tool to detect and avoid overcorrection in batch effect pipelines.

### 3.3. Application of the workflow in the comparison of two batch correction methods

#### ➢ Presentation of the use case

Two batch effect correction methods were evaluated for this use case, namely the LOESS-based (Dunn et al., 2011) and the ComBat correction (Johnson et al., 2012). While the former is one of the most widely used QC-based approaches in metabolomics, the latter provides a data-driven correction without the need of QCs. Based on the QC samples, LOESS involves the use of a local regression to adjust a model of the drift, which is then applied for data correction. It has the advantages of considering potential non-linearity. On the other hand, the ComBat method includes the use of empirical Bayes methods and prior knowledge (when available) to adjust a noise free model using linear regression (Johnson et al., 2012).

#### ➢ Dataset used for the application

The goals of this use case were to assess the ability of the workflow to capture different dimensions of the batch effect and evaluate its correction in a context of large-scale metabolomics. Moreover, this use case aimed at comparing the different batch correction methods in order to show a practical way and guide the user in the decision making when comparing different methods. The use case was conducted using QC samples from the predictive metabolomics study of neurological and metabolic toxicities induced by breast cancer treatment (Piffoux et al., 2024). Briefly, this dataset was obtained from serum samples analyzed using ultra-high performance liquid chromatography coupled with high-resolution Quadrupole Time-of-Flight (QToF) mass spectrometry (UHPLC-HRMS). A set of 992 serum samples was analyzed through 10 randomized batches involving 100 quality controls (QCs) to minimize technical biases. Almost 1,800 features were extracted and included in the current study. Full details of sample preparation and analysis are available in Piffoux et al, 2024 (Piffoux et al., 2024).

## Conclusion

In large-scale metabolomics studies, batch effects pose a significant challenge to data comparability and reliability. This work presents a workflow designed to assess the effectiveness and reliability of batch correction methods. By integrating multivariate analysis (PCA, clustering), numerical indicators (inter-batch and intra-batch dispersion, Coefficient of Variation, Isotopic Ratio), and univariate evaluation (using landmark compounds), the workflow provides a structured approach for determining the extent to which correction methods such as LOESS and ComBat help to improve data harmonization while preserving biochemical information. The proposed framework allows researchers to go beyond visual assessments, incorporating quantitative metrics to systematically compare correction methods, while ensuring that batch-wise variations are adequately mitigated without introducing new biases. By establishing this workflow, this study aims to standardize batch effect assessments, facilitating more reproducible and reliable metabolomic analyses.

The results obtained using the proposed workflow highlighted key differences between LOESS and ComBat batch correction methods, offering valuable insights for selecting and evaluating these approaches. Multivariate analyses (PCA and HCA) demonstrated that both methods effectively reduced batch effects, but the LOESS correction introduced more outliers, while achieving stronger batch-wise alignment. Numerical assessments, including inter-batch dispersion and coefficient of variation, confirmed that they both significantly improved data comparability, with LOESS providing better intra-batch consistency but slightly altering data structures. Univariate assessments on landmark compounds revealed that ComBat better preserved isotopic ratios integrity, whereas LOESS reduced variation more strongly. These findings underline the importance of selecting correction methods based on study objectives. LOESS could be recommended when the primary goal is minimizing batch effects at the risk of structural modifications, while ComBat is preferable for preserving intrinsic data properties. By following this workflow, users can systematically assess the reliability of batch corrections, ensuring both statistical accuracy and biological relevance in metabolomics studies.

## Data and code availability

All data and codes used in this work are available in a Zenodo archive (https://doi.org/10.5281/zenodo.16569697) and the methods will soon be available in Workflow 4 Metabolomics (16).

## Supporting information

**Suppl Figure 1:** Bivariate dispersion visualizations for different reference compounds, using QCs analyzed within nine batches on a large-scale study.

## Acknowledgments

All metabolomics analyses were funded and performed within the MetaboHUB French infrastructure (ANR-11-INBS-0010). E. Salanon is recipient of a doctoral fellowship from the INRAE DIGIT BIO metaprogramme and Geneva University.

## Author Contributions

Conceptualization: Blandine Comte, Estelle Pujos-Guillot, Julien Boccard,

Data curation: Delphine Centeno and Stéphanie Durand

Methodology: Elfried Salanon and Estelle Pujos-Guillot, Julien Boccard,

Resources: Estelle Pujos-Guillot, Julien Boccard,

Code/Software: Elfried Salanon

Supervision: Blandine Comte, Estelle Pujos-Guillot, Julien Boccard,

Writing – original draft: Elfried Salomon

Writing – review & editing: all authors

